# Computational modelling of the suppression of optic nerve fibre

**DOI:** 10.1101/2025.01.27.634990

**Authors:** Ariastity Pratiwi, Orsolya Kekesi, Alejandro Barriga-Rivera, Gregg Suaning

## Abstract

Retinal neuroprostheses aim to restore vision in degenerated retina but face challenges with low selectivity, where electric current activates both intended and neighbouring neurons. It was hypothesized that optic nerve suppression could improve selectivity using frequency-induced neuromodulation (FIN) with sinusoidal waveforms to selectively block transmission based on optic nerve fibre diameter. To explore the electrical parameters for diameter-based selective suppression, computational models of retinal ganglion cells (RGCs) and the optic nerve were given sinusoidal currents between 25-10000 Hz. FIN was found to induce complete conduction block, with the highest probability of suppression at frequencies above 500 Hz. Additionally, axon fibre diameter influenced the frequency and amplitude of FIN that inhibited conduction, with the highest selectivity between ON and OFF RGC fibres occurring at a diameter of 0.5 µm. Nodal sodium channels were involved in inhibition at frequencies above 500 Hz, supporting findings from studies of peripheral nerve fibres. Overall, suppressing a subset of optic nerve fibres could potentially enhance the selectivity of a bionic eye system by filtering out confounding visual information. However, practical application may be hindered by variability in RGC morphology and biophysical properties, and the possibility of filtering intended stimuli.

**Author summary:** Retinal neuroprostheses deliver electrical stimulation to retinal cells to produce perceptions of vision called phosphenes. However, the lack of selectivity associated with electrical currents leads to unnatural cell activations, which in turn limits the usability of many visual prostheses. To improve selectivity, we proposed a novel method that involves suppressing the conduction of action potentials along a subset of axon fibres, which carry the information from the eye to the brain’s visual centre. We believe that this could remove superfluous information caused by electrical stimulations. Our findings demonstrated that sinusoidal electrical currents can be given to the optic nerve fibres to induce suppression, and that the effectiveness of suppression is influenced by axon fibre diameters. This suggests that modulating the response of the optic nerve can lead to selective suppression based on the axon fibre diameters, and consequently improving the performance of a retinal neuroprosthesis.

## Introduction

Visual neuroprostheses use electrical currents to elicit responses from surviving neural cells to create *phosphenes* – visual sensations that ideally appear as distinct and localised perceptions of light. Ongoing research focuses on improving stimulation efficacy to produce selective activation, *i.e.,* only activating a small subset of neurons with a specific function at the intended location. However, the complexity of the retina, with millions of neurons serving various functions, presented an incompatibility with electrical current stimulations. Lateral spreading of the current can cause unintended activation of RGCs in addition to the intended cells to be stimulated. As neighbouring RGCs may codify different visual information [1], unintended activations can increase the complexity of phosphenes, compromising the benefits of the prosthetic system.

Strategies to improve stimulation selectivity include electrical pulse optimisation at the retina to target a specific neuron type [2]. Instead of attempting selective stimulations within the retina, we propose that neuromodulation at a secondary site - the optic nerve - can inhibit (block) some neural activation, which we posit can be useful in reducing unwanted activations. The idea of simultaneous modulation was proposed by Guo *et al.* [3] and this study explores the idea further. This approach could limit the information that reaches the visual centres in the brain by blocking a subset of axons. Conveniently, optic nerve anatomy enables the application of this approach to the full visual field [4].

Mammalian optic nerves, including human optic nerves, contain axon fibres of different diameters [5–7]. This study investigates selective blocking of spike conduction in optic nerve fibres of different diameters. As spatial segregation of different fibre size groups in the optic tract has been reported in different species [5, 8–9], this may suggest that fibre size-based blocking in the optic nerve would block neurons with different terminal locations.

Frequency-induced neuromodulation (FIN) of neural transmission has been shown in the peripheral and central nervous systems to trigger both activation and inhibition 10-13]. In the central nervous system, it was reported that reversible conduction blocking occurred at frequencies > 150 Hz [12], while 25-150 Hz was the frequency used for activations of retinal cells [13] and hippocampal cells [14]. Computational modelling studies indicated that fibre diameter affects the threshold amplitude required for suppression in the peripheral nervous system [15–17].

While the inhibition of peripheral nerve fibres is relatively prevalent in the academic literature, the inhibition of optic nerve conduction is largely unexplored. A notable exception is a study investigating the suppression of cat’s optic nerve conduction using mechanical pressure [18]. Here, Y-RGCs, associated with large-diameter optic nerve fibres, were suppressed for at least up to 4 days after the application of pressure [18], while the response carried by smaller fibres were left unchanged. Thus, it was concluded that mechanical pressure selectively inhibited the conduction of larger optic nerve fibres. The mechanism responsible for the inhibition was most likely myelin degeneration that could be retained for weeks [19]. As the suppression is not quickly reversible, the method using pressure would be less suitable than the application of electrical current to more rapidly produce reversible blockade.

Here, the feasibility of selective suppression according to fibre diameter and, by morphological association, the suppression of certain retinal ganglion cells (RGC) types using the application of sinusoidal electrical current to the optic nerve was explored using an *in silico* model.

## Methods

The computational model was developed following the two-stage modelling procedure as described by McNeal [20]. A finite element model (FEM) of retinal tissue and optic nerve based on rat’s anatomy was used to calculate the electric potential distribution across the tissue in response to electrical current injection. Then, the electric potential values were applied as extracellular activity to RGC models. Two types of RGCs, namely the ON and OFF RGCs, were investigated. RGC ion channel dynamics in both the soma and optic nerve axon fibre were defined using Hodgkin-Huxley-style equations [21] to determine the firing activity of the neuron. For a comprehensive overview and demonstration of the procedure described here, the reader is referred to our previous work on this topic [22].

### The anatomy of the retina and the optic nerve

COMSOL Multiphysics® 5.6 finite element modelling (FEM) was used, implementing the ‘Electric Current’ physics. A spherical cut-out of the eye globe was included in the model (Fig. 1A) with a single electrical conductivity value to represent the retina [23]. The optic nerve geometry was modelled with multiple layers [24]. A pair of penetrating electrodes with a conductive region at each tip (2000 µm^2^ of conductive area per electrode) in bipolar configuration were incorporated to deliver FIN. This configuration focuses the FIN modulation to a single optic nerve fibre node [25]. The rostral electrode (‘interference electrode’) delivered the FIN in reference to the caudal electrode (‘reference electrode’). The length of the optic nerve model was 5 mm to provide sufficient distance between the current source and the outer boundaries and avoid distortion in the electric potential calculation [26].

**Fig 1:**
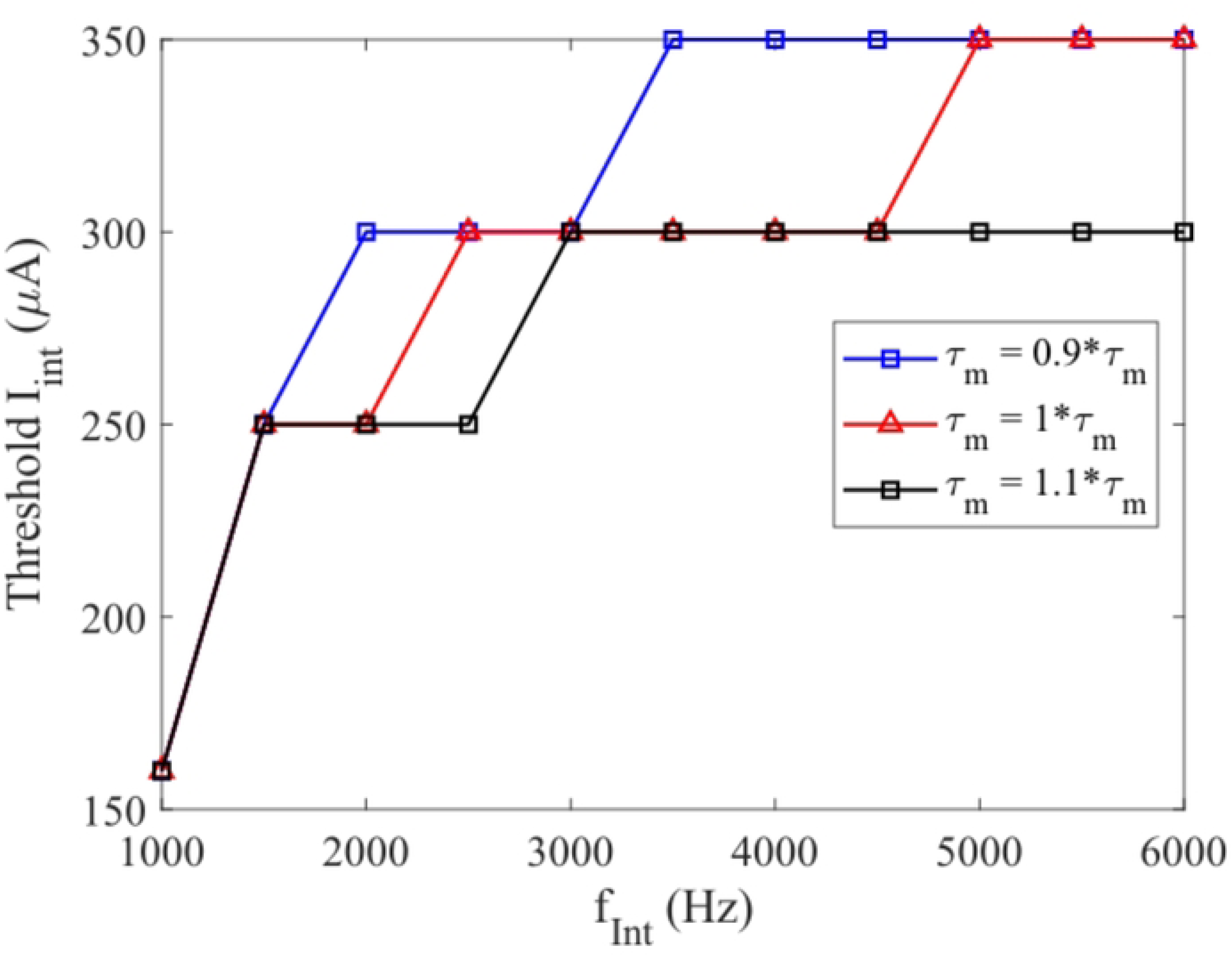
Geometry of the models. (A) The three-dimensional geometry of the eye and the optic nerve’s anatomy. The layers of the optic nerve (top right) include the fat, dura mater, CSF, pia mater, and the nerve fibre space. Bipolar FIN electrodes were geometrically modelled. (B) The division of the optic nerve fibre model. AH = axonal hillock, AIS = axon initial segment, DA = distal axon, MAF = myelinated axon fibre. H denotes the vertical length of the axonal section that distinguishes ON from OFF RGC. (C) The morphology of the RGC adapted from rabbit’s OFF RGC to fit rat’s alpha ganglion cells.

The choice of the geometry was based on rats as the computational model was intended to guide an *in vivo* study. Three different fibre diameters (d_F_) of 0.5, 2, and 3 µm, were used to represent rat’s nerve fibre population. The location of the electrode relative to the cell body differed for each nerve fibre diameter, with distances between the centre of the soma and the interference electrode being 1.27 mm, 1.38 mm, and 1.35 mm for d_F_ = 0.5, 2, and 3 µm, respectively. This configuration placed the cell body 270-380 µm from the optic disk, which is 1.1-1.6x the mean retinal thickness of wild-type Long Evans rats [27]. The reference electrode was placed 1 mm away from the interference electrode

To imitate action potential spikes produced by retinal stimulations, the RGC’s cell body was given an intracellular current injection, avoiding the superposition (crosstalk) effect of two distinct extracellular current injection sites. In practice, isolation of the retinal and optic nerve stimulations may require separate electrical circuits for each site, and/or local reference electrodes to limit the spatial spreading of electrical current. The geometrical and electrical parameters used are listed in S1 Table and were based on various experimental findings [28–31].

### RGC model

The morphologies of the ON and OFF RGC models, extending from the soma and the distal, nonmyelinated section of the axon (from soma to DA in *Fig. 1B*), were adapted from mouse OFF RGC model retrieved from Neuromorpho.org [32–34]. The dendritic field diameter was rescaled by a factor of 0.3 from the original to 400 μm to model rat’s α-RGC [35]. The axon was divided into the axon hillock (AH), a sodium-rich Axon Initial Segment (AIS), and a distal axon (DA) segment [36] (Fig.1B- C). The length and diameter of each section of the neuron are shown in S2 Table. The ON and OFF RGC models shared the same morphology, but the vertical length of the axonal section between the AH and soma of the OFF RGC was longer by 40 μm than the ON RGC’s [37]. While the kinetic equations were the same, the distribution of the ion channels differed between the ON and OFF RGC models (Table I). The kinetic equations followed that of Guo et al’s model [36], which modified Fohlmeister’s model of tiger salamander retina to include the Ca_T_ and HCN channels [38]. All membranes in the RGC model were assigned the same values for axial resistivity and membrane conductance, which are 0.11 kΩ.cm and 0.259 mS.cm^−2^ respectively [36].

**Table 1:** The modified conductance values for On and Off RGC models. D = Dendrite, S = Soma, AH = Axon Hillock, AIS = Axon Initial Segment, DA = Distal 131 Axon, in mS.cm-2

|  |  | **D** | **S** | **AH** | **AIS** | **DA** |
| --- | --- | --- | --- | --- | --- | --- |
| **ON RGC** | $g_{\text{Na}}$ | 50.2 | 112.7 | 94.1 | 220.9 | 110.8 |
| | $g_{\text{K,dr}}$ | 41.3 | 39.5 | 33.8 | 43.2 | 67.5 |
| | $g_{\text{K,Ca}}$ | 0 | 3.27 | 4.02 | 4.23 | 2.94 |
| **OFF RGC** | $g_{\text{Na}}$ | 38.7 | 128.6 | 71 | 201.9 | 80.5 |
| | $g_{\text{K,dr}}$ | 41.3 | 39.5 | 33.8 | 43.2 | 67.5 |
| | $g_{\text{K,Ca}}$ | 0 | 3.27 | 4.02 | 4.23 | 2.94 |

The conductance values of the sodium (g_Na_), delayed rectifier potassium (g_Kdr_), and calcium-activated potassium (g_Kca_) channels, were optimised to replicate the behaviour of ON and OFF RGCs when given sinusoidal stimuli reported in literature [2, 39]. The optimisation process involved a Markov Chain Monte Carlo (MCMC) method [36] with a Metropolis-Hastings algorithm [41–42]. The conductance values were iteratively updated to minimise a loss function. The following constraints were followed to produce a realistic model: g_Na_ is the highest at the AIS [43–44], g_Na_ at the AIS was between 100-500 mS/cm^2^, and g_Kdr_ in all regions between 32-150 mS/cm^2^ [45]. g_KCa_ values were freely changed. The optimisation results are presented in S1-S2 Files.

### Optic nerve model

As the extraocular portion of the optic nerve fibres are myelinated in most mammals, including humans [24, 46], only myelinated optic nerve fibres were modelled. Accordingly, the model only considers ion channels that have been identified at the nodal membranes of the optic nerve fibre, as most ion transfer happens at the nodes of Ranvier. McIntyre, Richardson and Grill (MRG) double cable model was followed [47]. The nerve fibre, except for the node, was surrounded by myelin. Each optic nerve fibre was segmented into different regions (Fig. 2), and the myelin thickness, as well as the segment length increases as the inner diameter increases. The ion channel kinetics were taken from the mammalian optic nerve fibre model [29, 48].

**Fig 2:**
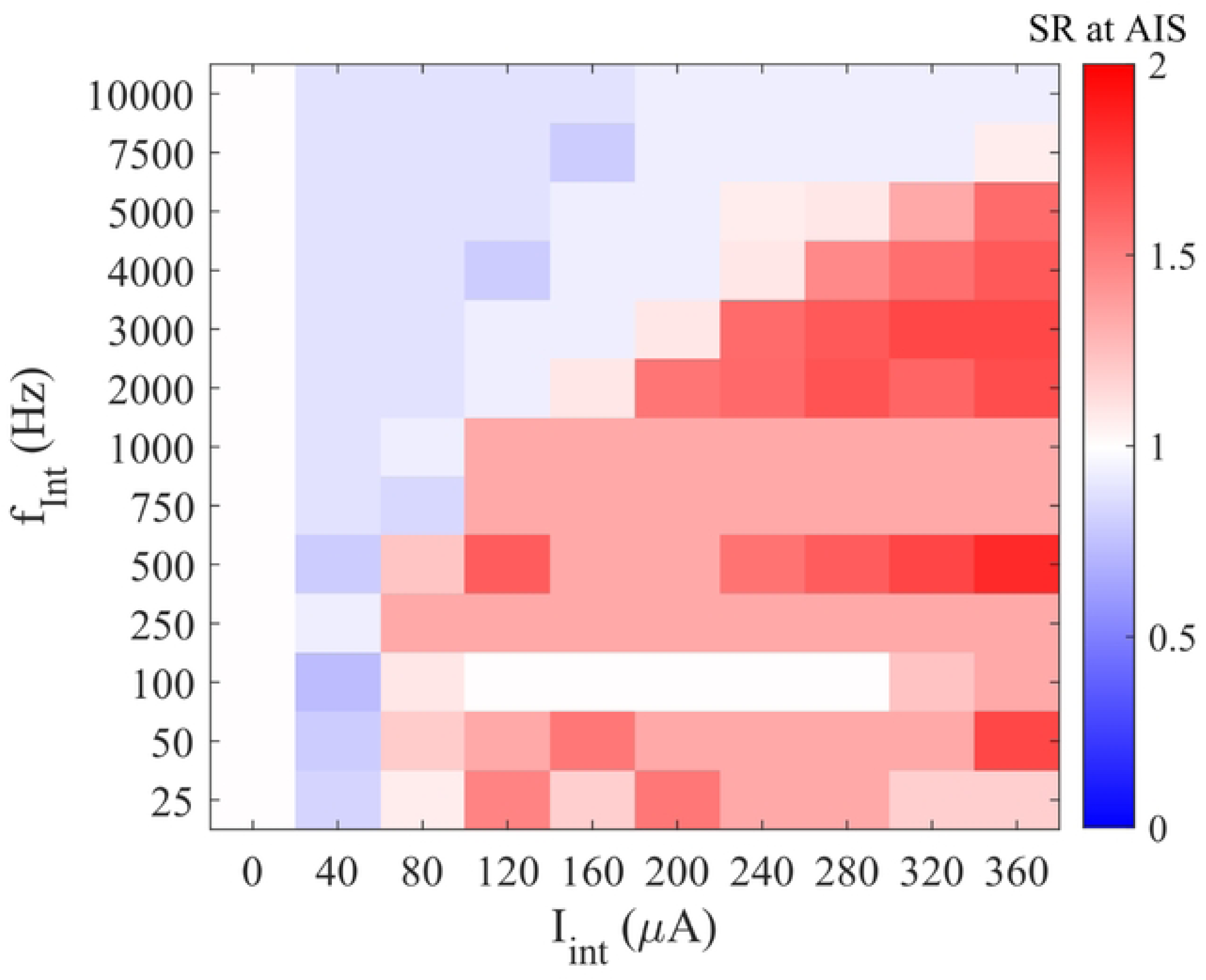
The division of optic nerve fibre morphology. N = nodal, PN = paranodal, JN = juxtaparanodal, and IN = internodal compartments. Each node-to-node unit contained 2 PNs, 2 JNs, and 6 INs. D1 and D2 are the nodal diameter and the fibre diameter, respectively. L denotes the length of each compartment.

As the original MRG model did not include fibre diameters that are representative of mammalian optic nerve fibres, which is below 2 µm [5–6], extrapolated geometrical parameters from [45] were used. According to Reese, the peaks of the fibre spectrum occurred around 0.75 µm and 1.5 µm, represented by d_F_ = 0.5 µm and 2 µm. A larger diameter of d_F_ = 3 µm represented coarse fibres that more commonly found in the central portion of the optic nerve in rats [5]. The model was validated by comparing the conduction velocity of the models to experimental findings in S3 File.

### Stimulation parameters

An intracellular current injection pulse (square, monophasic) inside the RGC’s soma at a threshold level for 40 ms (termed ‘activating stimulus’, I_stim_). The current used was just sufficient to produce a consistent spike train during the simulation period. At the same time, extracellular FIN was delivered via the interference electrode at the optic nerve. FIN amplitude (I_int_) ranged from 0-360 µA in steps of 5 μA, and the frequency (f_int_) ranged from 25-10,000 Hz. This frequency range covered the frequencies which have been shown *in vitro* to produce suppression in the CNS (0.5-150 Hz), as well as the kHz range that were shown *in vitro* to produce differential response between ON and OFF RGCs [2, 12, 49].

The simulation was run for 50 ms after the onset of the I_stim_ pulse, and the number of action potential spikes within this period were counted. Only the spikes whose peak magnitude was above 0 mV were included in the analysis. A suppression was said to have occurred if the number of action potential spikes detected at the probing point of the optic nerve fibre (the most distal node) was less than the initial number of spikes at threshold amplitude of I_stim_ [11]. The initial number of spikes was measured at the probing point when the cell body was given the activating stimulus I_stim_ without the interference (i.e., I_int_ = 0).

## Results

### Suppression of optic nerve conduction was influenced by the fibre diameter

After model optimisation, the responses of the RGCs’ optic nerve fibres to FIN were evaluated at the nodal region. The extent of suppression was quantified by the spike ratio (SR), defined as:

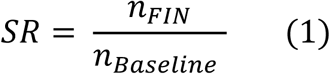

where *n*_FIN_ is the number of spikes produced when I_stim_ was delivered simultaneously with FIN at a given combination of I_int_ and f_Int_, while *n*_Baseline_ is the number of spikes produced when I_stim_ was delivered alone. Hence, an SR value of 1 (e.g., the column of I_Int_ = 0 μA in Fig. 3) indicates no suppression, while SR<1 indicates a reduction in the spike number (suppression) and SR>1 indicates an increase in the spike number (excitation).

**Fig 3:**
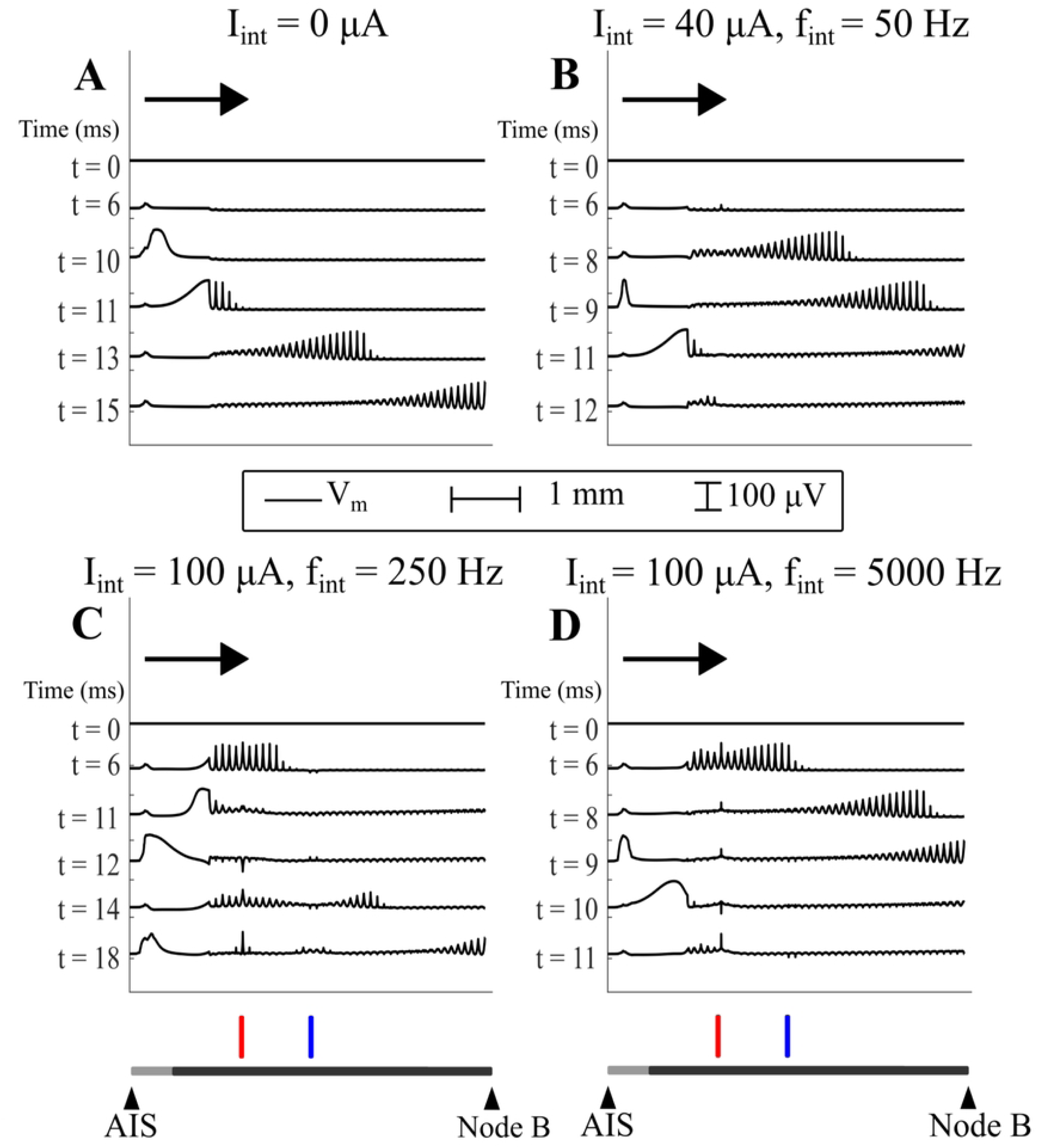
Effect of FIN varies based on the sinusoidal frequency and amplitude. (A, B) Colour grids of SR produced by combinations of IInt and fInt for ON and OFF fibres at different dF. IInt = 0 column indicated the SR when only Istim was delivered, hence the value was always 1. (C) The ratio between SR of ON and OFF RGC fibres. Black squares showed that the FIN parameters suppressed ON fibres more efficiently than the OFF fibres, gray squares showed no difference in the SR between ON and OFF fibres, and the white squares showed that the FIN parameters suppressed OFF fibres more efficiently than the ON fibres. Red squares indicate the FIN parameters that caused excitations in both ON and OFF fibres and thus are not appropriate for FIN application.

Both fibre diameter and RGC type gave rise to a variation in the SR produced by some combinations of f_Int_ and I_int_. Suppression was produced at all f_Int_, but reappearance of spikes occurred, making it difficult to identify a threshold I_int_ for suppression at most f_Int_. This was exemplified by f_Int_ > 750 Hz for d_F_ = 3 µm, where lower I_int_ produced SR close to 0, while increasing the I_int_ produced excitation.

The extent of FIN suppression between ON and OFF fibres was compared by dividing the SR values of the ON fibre by that of OFF fibre for each combination of f_Int_ and I_int_ (Fig. 3c). At d_F_ = 0.5 µm, most FIN parameters were more successful in suppressing ON fibres. At d_F_ = 2 µm, there was no apparent difference, while at d_F_ = 3 µm, there was a delineated area in the grid that more effectively suppressed ON fibres at I_int_ = 70 - 260 µA and f_Int_ = 750 – 10000 Hz, adjacent to an area that more effectively suppressed OFF fibres at I_int_ = 100 - 360 µA and f_Int_ = 750 – 3000 Hz. When results from all d_F_ of ON and OFF fibres were considered, almost all f_Int_ frequencies showed caused unintended excitation at some amplitude, apart from f_Int_ = 25 – 100 Hz.

Further, an index called suppression probability per f_Int_ was calculated by summing the SR values across all I_int_ and taking its inverse, then normalised to the maximum probability found for each fibre diameter (Fig. 4a) or each fibre type (Fig. 4b). All fibres demonstrated three separate frequency domains that may indicate different types of suppressions, which are (i) low frequency (25 - 250 Hz), (ii) intermediate frequency (500 - 1000 Hz), and (iii) high frequency (2000 - 10000 Hz). Suppression probability tended to decrease with frequency for low frequency, increase with frequency for intermediate frequency, and decrease with frequency for high frequency. The lowest suppression probability was consistently found at 250 Hz for all the fibre model, while the highest suppression probability occurred at various frequencies, including the lowest f_Int_ tested of 25 Hz, as well as around 750-2000 Hz. The difference between the suppression probability for between ON and OFF fibres was the most consistent at d_F_ = 0.5 µm, with the largest difference at the high frequency domain. Furthermore, the maximum separation between diameters was achieved at high frequency suppression for both ON and OFF fibres. The results suggested that high frequency was the most effective in producing selective suppression in optic nerve fibres based on fibre diameters, as well as the RGC type. This demonstrates the possibility of exploiting the differences in characteristics of ON and OFF RGCs in producing selective suppression. However, this observation is only limited to one fibre diameter (d_F_ = 0.5 μm).

**Fig 4:**
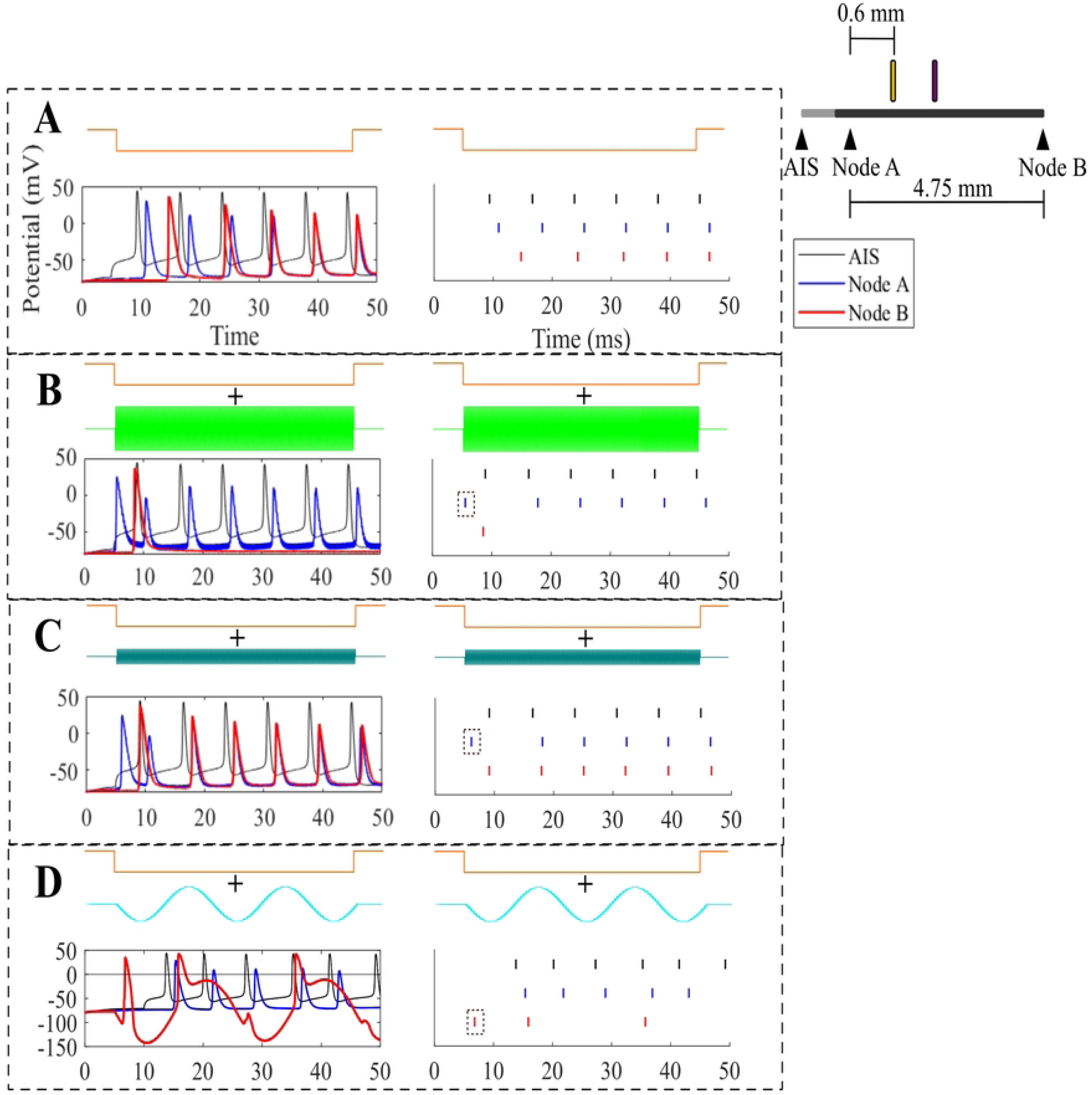
Probabilities of suppression grouped by fibre diameter and type of RGC. Normalised probability of suppression against the frequency of FIN for ON and OFF fibre models grouped by (A) the fibre diameter and (B) the RGC type

### The time course of different mechanisms of FIN suppression

To identify the contributions of FIN on action potential conduction, transmembrane potentials across the whole cell for ON fibre (d_F_ = 0.5 μm) were observed. The number of spikes detected at three different points (AIS, node A and node B), were summarised in raster plots (Fig. 5). Node A was located 0.6 mm proximal to the interference electrode, and node B was located 4.2 mm distal to the interference electrode. Node A reflects the effect of FIN on the most proximal node, while node B reflects the spikes delivered to the brain.

**Fig 5:**
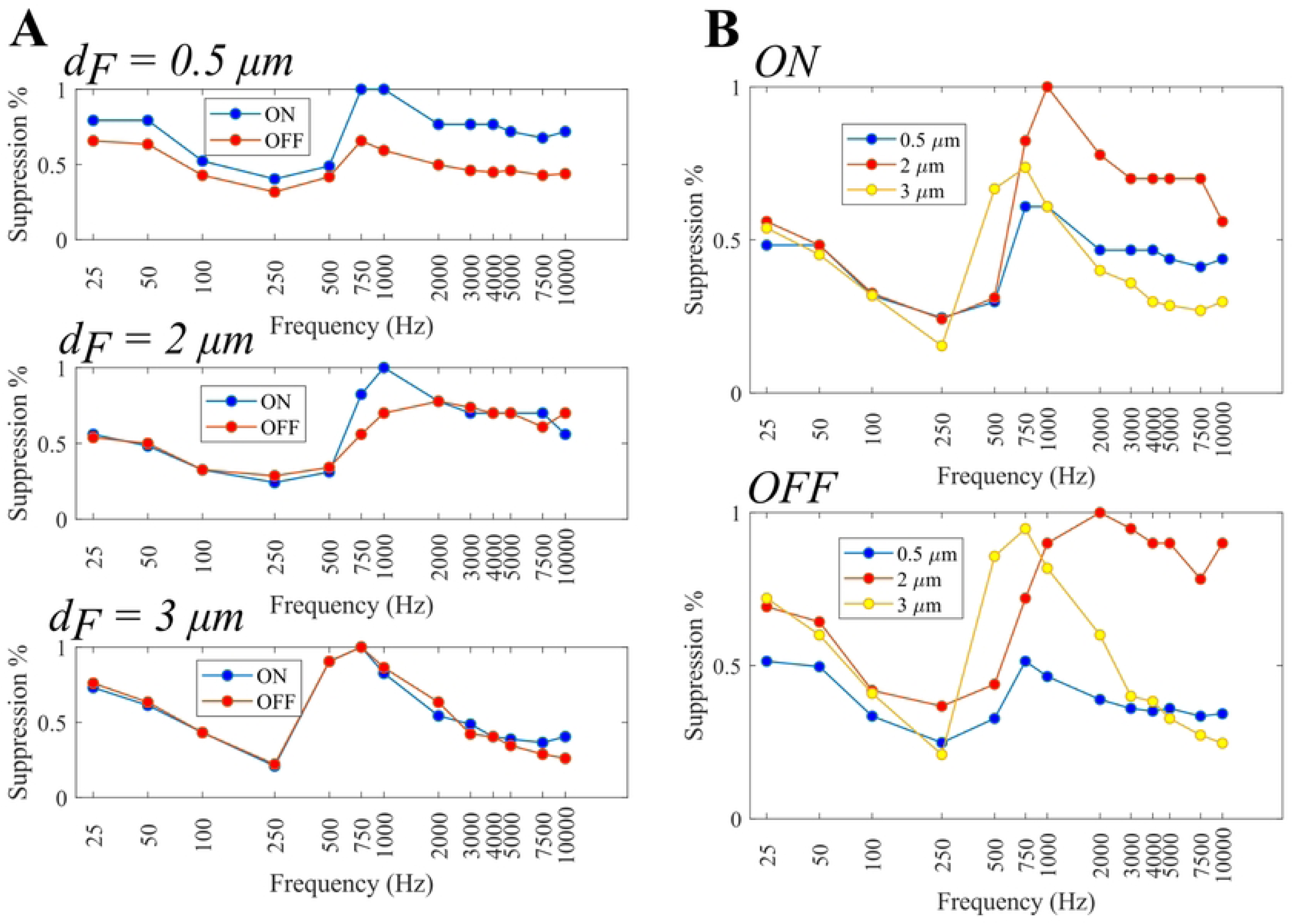
Different responses of optic nerve fibres to FIN. (A) Only Istim was delivered. All spikes originated from the AIS travelled to node B. (B) Istim was delivered together with FIN (Iint = 300 µA, fInt = 2000 Hz). Complete suppression occurred, and only one spike was detected at node B. (C) Istim delivered together with FIN (Iint = 120 µA, fInt = 2000 Hz). Excitation occurred as FIN triggered an additional action potential spike in the optic nerve while still allowing the spikes generated at the AIS to conduct to node B. (D) Istim was delivered together with FIN (Iint = 40 µA, fInt = 50 Hz). FIN elicited an action potential spike in the optic nerve, but at the same time suppressing some of the action potentials generated at the AIS, resulted in a partial suppression.

Three effects of combining I_stim_ with FIN were observed: complete suppression, excitation, and partial suppression. Complete suppression occurred when FIN blocked all the spikes from the soma. When only I_stim_ was delivered, a train of action potentials generated at the RGC travelled along the optic nerve, shown by the slight delay in the spike timings at Nodes A and B (Fig. 5A). Using f_Int_ = 2000 Hz and I_int_ = 300 µA, complete suppression occurred and all the RGC spikes were blocked (Fig. 5B). However, a single spike was generated at the optic nerve whose timing preceded the first RGC spike. Note that in all cases of complete suppression, there was always one spike at Node B. Complete suppression mostly happened at the high and intermediate frequency domains. This suppression did not happen when the charge was kept the same but using lower f_Int_ and I_int_, e.g., f_Int_ = 250 Hz and I_int_ = 35 µA (result not shown). This points to the existence of a threshold amplitude where this type of suppression could happen, and thus this effect being frequency- and amplitude-dependent, rather than merely charge-dependent.

Next, FIN could cause excitation that resulted in the increase of SR (Fig. 5C). Lowering I_int_ to 120 µA at f_Int_ = 2000 Hz generated additional action potential spikes at the optic nerve fibre without blocking the action potentials produced at soma. In partial suppression, FIN blocked the RGC spikes, but at the same time generated action potentials at the FIN site. This likely involved a mix of hyperpolarisation- induced block and suppression via antidromic passage of spikes. Partial suppression is demonstrated in Fig. 5D (f_Int_ = 50 Hz and I_int_ = 40 µA). All the spikes from the cell body travelled to node A, but the hyperpolarisation from FIN caused some of the spikes at node B to be subthreshold, as seen at t = 27 ms. However, additional spikes were generated by FIN itself, signified by the onset of the first spike at node B that preceded the first spike detected at the AIS. The hyperpolarisation-induced block, or commonly called anodal block, is consistent with the finding that this partial suppression occurred mostly at the low frequency domain with a probability of suppression that decreases with increasing f_Int_. This type of block is thus charge dependent.

### The proximity of the FIN electrode and AIS caused secondary suppression and the differential response between ON and OFF fibres

The application of FIN to the nodal membrane could generate action potential spikes that are independent of the I_stim_, potentially causing secondary suppression via the antidromic passage of action potential spikes along the nerve fibre. When FIN was absent, action potentials were generated at the AIS at the beginning of I_stim_, and the spikes travelled towards the last node (Fig. 6A). Adding FIN at f_Int_ = 50 Hz and I_int_ = 40 µA depolarised the node below the interference electrode and generated additional spikes that traversed ortho- and antidromically towards the cell body and the distal node (Fig. 6B). Some of the spikes travelling antidromically towards the cell body caused a secondary suppression due to the refractory period of the RGC itself, reducing the number of spikes detected at the AIS compared to the baseline when FIN was absent (Fig. 5D). This type of suppression, however, was found together with the elicitation of spikes by FIN application at the membrane below the reference electrode. This is demonstrated by FIN at f_Int_ = 250 Hz and I_int_ = 100 µA (Fig. 6C) and f_Int_ = 5000 Hz, 100 µA (Fig. 6D).

**Fig 6:**
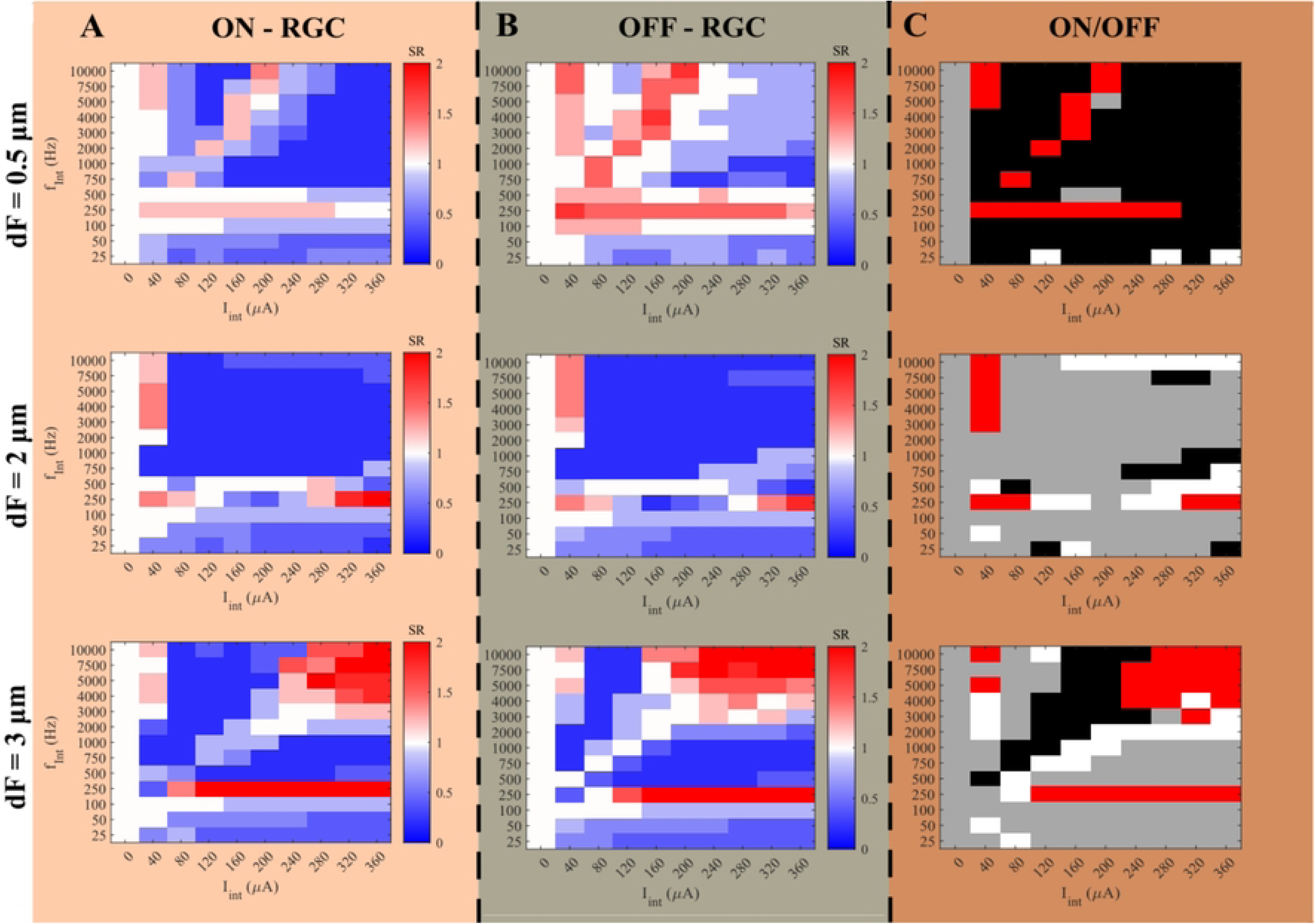
The propagation of the action potential spike with FIN at different time points. X-axis showing the location along the optic nerve fibre, aligned with the AIS on the left and the most distal node on the right (node B). Time in milliseconds where the event happened was indicated by t, where t = 0 is the stimulus onset. Arrows denote the orthodromic direction of spike propagation. (A) Only Istim was delivered, and all spikes detected at node B originated from the AIS. (B) IInt = 40 μA at fInt = 50 Hz was delivered together with Istim. Two locations of action potential generation were found; one at the AIS, and the other just below the interference electrode. Spikes from the membrane below the interference electrode travelled in two directions. (C) IInt = 100 μA at fInt = 250 Hz was delivered together with Istim. An additional location of action potential generation was created underneath the reference electrode. (D) IInt = 100 μA at fInt = 5000 Hz was delivered together with Istim. Two locations of action potential generation were found, same as (B).

Another consequence of FIN is the differential response found for ON and OFF fibre models. As the ON and OFF fibre models used the same formulations for the myelinated fibre section, the interaction between FIN and the cell body is the cause for any variation in the SR. This could be due to direct stimulation of the AIS by FIN, or through the interaction between the antidromic passage of action potential spikes from the fibre’s node and the action potential spikes from the AIS. At d_F_ = 0.5 μm, applying FIN at high frequency domain resulted in complete suppression in many instances for ON fibre, while it only caused partial suppression on the OFF fibre. Fig. 7 showed the ratio of spike counts between ON and OFF fibres (SR at AIS) at different combinations of f_Int_ and I_int_. The same combination of FIN parameters generated more spikes at the AIS and node A of the ON fibre compared to OFF fibre, which is likely to be caused by the higher concentration of the sodium channels at the AIS. This suggests that the suppression mechanism at high frequency domain was influenced by the membrane’s response upstream from the FIN site, and that a more active AIS and node A increases the probability of complete suppression.

**Fig 7:**
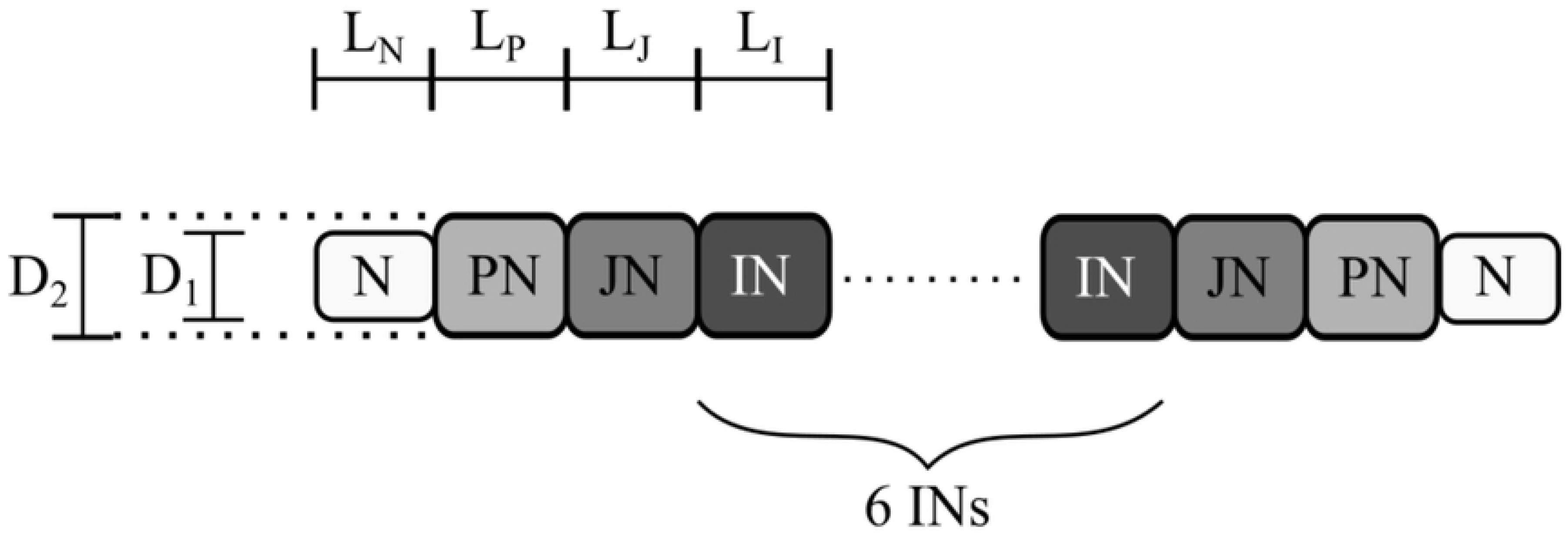
The ratio of the spike counts at AIS (SR at AIS) between ON fibre and OFF fibre at dF = 0.5 μm. The FIN domain where ON suppression was more successful than OFF suppression was predominated by higher SR at AIS.

### Sodium channel kinetics affecting the suppression at higher stimulus frequencies

To test that Na_f_ channels of the optic nerve fibre’s node contributed to the complete suppression at high frequency domains, the channel’s opening time constant (τ_m_) was varied and the effect on the block threshold amplitudes at different stimulus frequencies was observed. Changing the time constants of the Na_f_ channel changed the minimum frequency required to cause the high frequency block, as well as the amplitude thresholds. For the ON fibre model with d_F_ = 0.5 μm, the combinations of f_Int_ and threshold amplitude at the corresponding frequency for three different time constants were plotted (Fig. 8). Changing τ_m_ to a larger value tended to reduce the stimulus amplitude required to achieve high frequency blocking at a particular f_Int_, meaning that making the activation of sodium channels slower would facilitate high frequency blocking. The contribution of Na_f_’s time constant suggests that the inability of the sodium channels to open and close rapidly enough to follow the frequency of stimulus may cause the failure of spikes’ elicitation.

**Fig 8:**
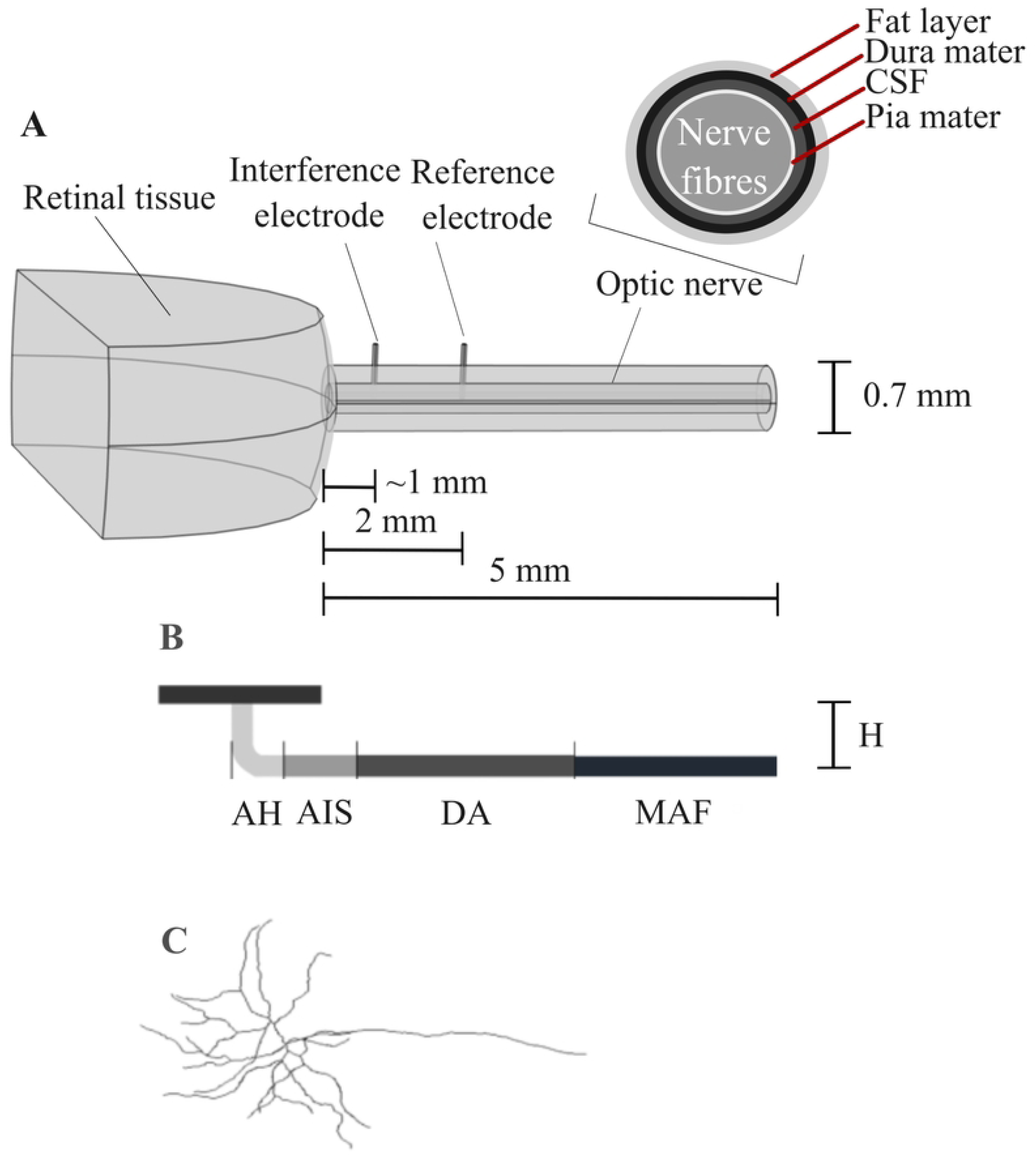
The threshold current required to induce a high frequency blocking for different τm values. The threshold current at the same fInt was reduced as the time constant τm was increased.

## Discussions

### Identification of different causes of suppression in the optic nerve

While conduction blocking of nerve fibres has been widely investigated, existing studies focused on the conduction block in peripheral nerve fibres, which are larger in size [50] and/or have different ion channel distributions [51]. For example, the nodal regions of the mammalian peripheral nerve fibres contain an abundance of slow potassium channels [52], unlike the optic nerve fibre model [49]. Hence, proposed mechanisms such as activation of potassium channels curtailing conduction [52] were unlikely to be the cause of suppression observed in this model.

The simulation revealed variations in SR values and FIN efficacy at different sinusoidal current frequencies, suggesting different mechanisms of suppression. The current model demonstrated a blocking mechanism where action potentials did not travel further due to hyperpolarisation near the interference electrode, consistent with anodal or cathodal block effects [16]. At higher FIN frequencies (f_Int_>500 Hz), another type of suppression was demonstrated, similar to that reported in peripheral nerve fibres [11, 17]. Artificially changing the time constant of the gating parameter (τ_m_) to slow down the opening/ closing of the sodium channels slower lowered the inhibition threshold at higher f_Int_. This points to the fast sodium channels contributing to the suppression, possibly incapable of following the rapid oscillation of the sinusoidal current. This agrees with the finding from the spectral analysis of sodium channel’s dynamics by Ackermann, *et al.* [10].

### Evidence of selective suppression based on axonal fibre diameter in the optic nerve

FIN was shown to suppress axonal conduction in the optic nerve, with differences in suppression efficacy based on fibre diameter, and to a lesser extent, RGC subtype. Unlike in peripheral nerve models [17], the correlation between the suppressive f_Int_ and I_Int_ combinations and the fibre’s diameter was less prominent in the optic nerve. It was expected that the suppression threshold would increase with fibre diameter, and the threshold changes more steeply at smaller fibre diameter [17, 53]. Such trends were not seen in the current results, mostly owing to the reappearance of spikes when the current amplitude was increased further, making it challenging to find a suppression threshold. This observation reflected that of vagus nerve fibres, whose diameters are close to the current optic nerve fibre models (1, 2, and 5.7 μm) [54]. These suggest that the relationship between fibre diameter and suppression threshold only holds for a certain range of fibre diameters. It is well-known that neural activations are influenced by morphological factors. In the optic nerve fibre model, the ratio between the nodal area (where the activation happens) and the resistive non-nodal area was higher than in the original MRG model, possibly lowering the likelihood of suppression and making the membrane easily activated. It should also be reiterated that the MRG model was validated using fibres large peripheral nerve fibres, and thus *in vitro* or *in vivo* validation involving smaller fibre diameters would be useful.

However, separating the FIN frequencies into three different ranges demonstrated that the difference in suppression between diameters was the most significant in the high frequency range for both ON and OFF fibres. This shows that the high frequency blocking mechanism is the most sensitive to fibre diameter. This finding is also in line with observations in peripheral nerve fibres, where consistent block could be achieved using higher frequency current at 3 kHz and above [10–11, 17].

### Suppression was modified by the dynamics of the RGCs, but this effect might be diminished in larger mammals

Including cell body morphology and ion channel kinetics in the nerve fibre model was shown to affect its response to FIN. Here, both ON and OFF fibre models used the same morphological formulation and ion channel kinetics at the myelinated axon. However, the responses at the AIS affected the nerve fibre’s response to FIN. This could be due to the electric field produced by FIN directly affecting the membrane close to the cell body or mediated by the secondary suppression caused by the antidromic passage of the spike generated at the FIN site. In these cases, the morphological and biophysical variations near cell body could modulate the cell response, resulting in the lower suppression probability of OFF fibres compared to ON fibres.

This result suggests the possibility of using FIN to produce selective suppression based on the morphological and biophysical variations of different RGC types. However, this finding is likely to be influenced by our model’s geometry that followed rat’s anatomy. Direct modulation was caused by the high conductivity of the optic nerve in the longitudinal direction, which was facilitated by the proximity of the FIN electrode and the cell body. It is possible that this effect would be minimal when applied in larger mammals or humans, due to the larger distance between the cell body of the RGC and the potential placement site of the FIN electrodes. Similarly, interaction between spikes generated at the AIS and spikes generated at the FIN site is less likely to happen in larger animals due to the decay of the action potential in space.

### Applications of selective suppression based on fibre size in a bionic eye system

This study looked at the possibility of suppressing action potential selectively based on the diameter of the optic nerve fibre. As humans’ optic nerves contain axons of various sizes, this method could be applicable to bionic eye systems. It could also contribute to functional selectivity. While a direct relationship between fibre size and cell function has not been completely established, correlations between fibre size groups and RGC functions, such as cats’ X, Y and W RGCs [55], have been shown. The correlation between functions and fibre diameter in other RGC types or other species remains to be investigated. For ON and OFF RGCs, differences in the cell body’s morphology and ion channel kinetics have been characterised [56–57], while variations in the myelinated axon region are not yet known.

An important drawback of FIN suppression is the possibility of blocking intended signals, which will be counterproductive. At the optic nerve, where the fibre size is distributed randomly, applying FIN to block one group of fibres would be less useful, as the blocked fibres can originate from any portion of the retina. Thus, fibre size-based modulation should be used with other techniques to increase specificity, such as current shaping techniques [58] to better target the optic nerve fibres. Alternatively, FIN suppression can be applied to the optic tract, where fibre size groups are more spatially segregated [5, 8–9].

Furthermore, at least one spike was detected at the start of FIN application in all cases. This could be caused by the gradual change in the extracellular potential. In practice, activation should be avoided by optimising the way that FIN is delivered. For example, the suppressive FIN could be initiated with a smaller amplitude to avoid activation and slowly increasing it to the required current to achieve suppression. Alternatively, FIN can be applied continuously as a tonic current injection, so that the additional spike will only be produced once at the start of the FIN application, and not at the subsequent instances of retinal stimulation.

### Limitations

An important consideration is that the model was formulated using data from different species. While the morphological parameters of the ON and OFF RGCs were fitted to rat, but used electrophysiological data from both rabbit [2] and rat [35]. This was done due to data availability and is a common practice in computational modelling, but ideally the model should be verified with dataset from the same species. Furthermore, this model only included the ion channels in the nodal regions, because ion channels at the other regions of the nerve fibres have not been extensively investigated or validated [43]. Refinements of the model that include details regarding the variation of ion channels contained by axons of different RGC subtypes, particularly at the MAF region, should be implemented in future studies.

## Conclusions

Restricting the information that reaches the brain in the case of retinal stimulation can be beneficial. The study showed that FIN using sinusoidal currents could suppress action potential transmission along the optic nerve. The frequencies and amplitudes of current where the suppression happened were influenced by the diameter of the nerve fibre, and to a lesser extent the morphological and biophysical properties of the RGC itself. Multiple modes of suppression were shown by the model, but the most successful suppression mostly happened at FIN frequencies higher than 500 Hz. This mode of suppression was influenced by the inherent properties of the optic nerve fibre’s node’s fast sodium channel, as its time constant influences the current amplitude at which suppression occurred. Another mode of suppression, that involved an antidromic passage of action potential from the site of FIN towards the AIS, was observed, and caused the variation in the extent of suppression between ON and OFF RGC fibres. However, it is acknowledged that this effect might be diminished in larger mammals, where the distance between the FIN site and the RGC cell body is larger. Additionally, at least one additional spike was always produced at the FIN site at any combination of frequency and amplitude tested. It is proposed that FIN is either introduced with a slowly increasing amplitude or used as a continuous input to minimise the likelihood of creating excitation. *In vitro* or *in vivo* works to verify the optic nerve model and the phenomenon of suppression are necessary to consolidate the findings of this study.

## Supporting Information

**S1 Table: The geometrical and electrical parameters of the retina and optic nerve tissue. S2 Table: The geometrical and electrical parameters of the optic nerve fibres.**

**S1 Text: Optimisation of the RGC models. S2 Text: Validation of the RGC models.**

**S3 Text: Validation of the optic nerve fibres’ conduction velocity.**

**S1 Fig: Validation of the OFF RGC models’ spike counts per period compared to *in vitro* findings.** The spikes per sinusoidal FIN period of OFF-BT and OFF-S in response to sinusoidal extracellular stimuli at different frequencies, measured in vitro by Twyford et al. [2], compared to the spikes per period of the modified OFF RGC model in response to sinusoidal extracellular stimuli at the same frequencies.

**S2 Fig: Validation of the ON and OFF RGC models’ spike frequency compared to *in vitro* findings**. The spiking frequency of ON and OFF RGC models, in response to extracellular sinusoidal currents at a constant amplitude (ON RGC at 70 μA and OFF RGC at 76 μA), showing an increase of the spiking response with stimulus frequency, followed by a decrease at higher frequencies tested. The spiking frequencies for OFF RGC were always lower than the ON RGC.

**S3 Fig: Conduction velocities of the optic nerve fibre models**. The resulting conduction velocity of the optic nerve fibre model at d_F_ = 0.5, 2 and 3 μm. The data are fitted to a linear function y=5.1x-1.9.

## References

1. Masland RH. The neuronal organization of the retina. Neuron. 2012;76(2):266–80.

2. Twyford P, Fried S. The retinal response to sinusoidal electrical stimulation. IEEE Trans Neural Syst Rehabil Eng. 2015;24(4):413–23.

3. Guo T, et al. Mimicking natural neural encoding through retinal electrostimulation. In: Proceedings of the 2017 8th International IEEE/EMBS Conference on Neural Engineering (NER); 2017.

4. Veraart C, et al. Pattern recognition with the optic nerve visual prosthesis. Artif Organs. 2003;27(11):996–1004.

5. Reese BE. The distribution of axons according to diameter in the optic nerve and optic tract of the rat. Neuroscience. 1987;22(3):1015–24.

6. Ogden TE, Miller RF. Studies of the optic nerve of the rhesus monkey: nerve fiber spectrum and physiological properties. Vision Res. 1966;6(9-10):485-IN2.

7. Reese BE, Ho KY. Axon diameter distributions across the monkey’s optic nerve. Neuroscience. 1988;27(1):205–14.

8. Guillery RW, Polley EH, Torrealba F. The arrangement of axons according to fiber diameter in the optic tract of the cat. J Neurosci. 1982;2(6):714–21.

9. Reese BE, Guillery RW. Distribution of axons according to diameter in the monkey’s optic tract. J Comp Neurol. 1987;260(3):453–60.

10. Ackermann DM, et al. Dynamics and sensitivity analysis of high-frequency conduction block. J Neural Eng. 2011;8(6):065007.

11. Kilgore KL, Bhadra N. High frequency mammalian nerve conduction block: simulations and experiments. In: Proceedings of the 2006 International Conference of the IEEE Engineering in Medicine and Biology Society; 2006.

12. Jensen AL, Durand DM. High frequency stimulation can block axonal conduction. Exp Neurol. 2009;220(1):57–70.

13. Benazzouz A, et al. Responses of substantia nigra pars reticulata and globus pallidus complex to high frequency stimulation of the subthalamic nucleus in rats: electrophysiological data. Neurosci Lett. 1995;189(2):77–80.

14. Freeman DK, et al. Selective activation of neuronal targets with sinusoidal electric stimulation. J Neurophysiol. 2010;104(5):2778–91.

15. Wang Z, et al. Sinusoidal stimulation on afferent fibers can selectively activate different types of neurons in rat hippocampus. In: Proceedings of the 2019 41st Annual International Conference of the IEEE Engineering in Medicine and Biology Society (EMBC); 2019.

16. Tai C, Roppolo JR, de Groat WC. Analysis of nerve conduction block induced by direct current. J Comput Neurosci. 2009;27:201–10.

17. Bhadra N, et al. Simulation of high-frequency sinusoidal electrical block of mammalian myelinated axons. J Comput Neurosci. 2007;22:313–26.

18. Burke W, Burne J, Martin P. Selective block of Y optic nerve fibres in the cat and the occurrence of inhibition in the lateral geniculate nucleus. J Physiol. 1985;364(1):81–92.

19. Seddon HJ. Three types of nerve injury. Brain. 1943;66(4):237–88.

20. McNeal DR. Analysis of a model for excitation of myelinated nerve. IEEE Trans Biomed Eng. 1976;(4):329–37.

21. Hodgkin AL, Huxley AF. The components of membrane conductance in the giant axon of Loligo. J Physiol. 1952;116(4):473–96.

22. Pratiwi A, Kekesi O, Suaning G. Computational modeling of retinal neurons for visual prosthesis research—fundamental approaches. J Vis Exp. 2022;(184):e63792.

23. Resatz S, Rattay F. A model for the electrically stimulated retina. Math Comput Model. 2004;10(2):93–106.

24. Fukui Y, et al. Quantitative study of the development of the optic nerve in rats reared in the dark during early postnatal life. J Anat. 1991;174:37–46.

25. Johnson MD, et al. Neuromodulation for brain disorders: challenges and opportunities. IEEE Trans Biomed Eng. 2013;60(3):610–24.

26. Raspopovic S, Capogrosso M, Micera S. A computational model for the stimulation of rat sciatic nerve using a transverse intrafascicular multichannel electrode. Unpublished.

27. Brown AM, et al. Axonal L-type Ca²⁺ channels and anoxic injury in rat CNS white matter. J Neurophysiol. 2001;85(2):900–11.

28. Manola L, Holsheimer J, Veltink PH, et al. Modelling motor cortex stimulation for chronic pain control: electrical potential field, activating functions and responses of simple nerve fibre models. Med Biol Eng Comput. 2005;43(3):335–343.

29. Oozeer M, et al. A model of the mammalian optic nerve fibre based on experimental data. Vision Res. 2006;46(16):2513–24.

30. Struijk JJ, Holsheimer J, van Veen BK, Boom HB. Paresthesia thresholds in spinal cord stimulation: a comparison of theoretical results with clinical data. IEEE Trans Rehabil Eng. 1993;1(2):101–108.

31. Lee S, Kim H, Park S, et al. In-vivo estimation of tissue electrical conductivities of a rabbit eye for precise simulation of electric field distributions during ocular iontophoresis. Int J Numer Method Biomed Eng. 2022;38(1):e3540.

32. Neuromorpho.org. Available from: www.neuromorpho.org. Accessed January 10, 2025.

33. Akram MA, et al. An open repository for single-cell reconstructions of the brain forest. Sci Data. 2018;5(1):1–12.

34. Werginz P, Raghuram V, Fried SI. Tailoring of the axon initial segment shapes the conversion of synaptic inputs into spiking output in OFF-α retinal ganglion cells. Sci Adv. 2020;6(37):eabb6642.

35. Yang JM, et al. Development of a novel knockout model of retinitis pigmentosa using Pde6b- knockout Long-Evans rats. Front Med. 2022;9:909182.

36. Guo T, et al. Insights from computational modelling: selective stimulation of retinal ganglion cells. Brain Human Body Model. 2021;2020:233–50.

37. Ammermüller J, Kolb H. The organization of the turtle inner retina. I. ON- and OFF-center pathways. J Comp Neurol. 1995;358(1):1–34.

38. Fohlmeister J, Miller R. Impulse encoding mechanisms of ganglion cells in the tiger salamander retina. J Neurophysiol. 1997;78(4):1935–47.

39. Hadjinicolaou AE, et al. Frequency responses of rat retinal ganglion cells. PLoS One. 2016;11(6):e0157676.

40. Wang YC, et al. Multimodal parameter spaces of a complex multi-channel neuron model. Front Syst Neurosci. 2022;16:999531.

41. Hastings WK. Monte Carlo sampling methods using Markov chains and their applications. 1970.

42. Metropolis N, et al. Equation of state calculations by fast computing machines. J Chem Phys. 1953;21(6):1087–92.

43. Jeng J, et al. The sodium channel band shapes the response to electric stimulation in retinal ganglion cells. J Neural Eng. 2011;8(3):036022.

44. Fried SI, et al. Axonal sodium-channel bands shape the response to electric stimulation in retinal ganglion cells. J Neurophysiol. 2009;101(4):1972–87.

45. Fohlmeister JF, Cohen ED, Newman EA. Mechanisms and distribution of ion channels in retinal ganglion cells: using temperature as an independent variable. J Neurophysiol. 2010;103(3):1357–74.

46. Miyake E, Imagawa T, Uehara M. Fine structure of the retino-optic nerve junction in dogs. J Vet Med Sci. 2005;66:1549–54.

47. McIntyre CC, Richardson AG, Grill WM. Modeling the excitability of mammalian nerve fibers: influence of afterpotentials on the recovery cycle. J Neurophysiol. 2002;87(2):995–1006.

48. Li M, et al. A simulation of current focusing and steering with penetrating optic nerve electrodes. J Neural Eng. 2013;10(6):066007.

49. Muralidharan M, et al. Neural activity of functionally different retinal ganglion cells can be robustly modulated by high-rate electrical pulse trains. J Neural Eng. 2020;17(4):045013.

50. Ritchie JM. On the relation between fibre diameter and conduction velocity in myelinated nerve fibres. Proc R Soc Lond B Biol Sci. 1982;217(1206):29-35.

51. Wang H, et al. Heteromultimeric K+ channels in terminal and juxtaparanodal regions of neurons. Nature. 1993;365(6441):75-9.

52. Liu H, et al. The role of slow potassium current in nerve conduction block induced by high-frequency biphasic electrical current. IEEE Trans Biomed Eng. 2008;56(1):137–46.

53. Danner SM, Wenger C, Rattay F. Electrical Stimulation of Myelinated Axons: An Interactive Tutorial Supported by Computer Simulation. Saarbrücken, Germany: VDM; 2011.

54. Pelot NA, Behrend CE, Grill WM. Modeling the response of small myelinated axons in a compound nerve to kilohertz frequency signals. J Neural Eng. 2017;14(4):046022.

55. Williams RW, Chalupa LM. An analysis of axon caliber within the optic nerve of the cat: evidence of size groupings and regional organization. J Neurosci. 1983;3(8):1554–64.

56. Margolis DJ, et al. Dendritic calcium signaling in ON and OFF mouse retinal ganglion cells. J Neurosci. 2010;30(21):7127–38.

57. Werginz P, Raghuram V, Fried SI. The relationship between morphological properties and thresholds to extracellular electric stimulation in α RGCs. J Neural Eng. 2020;17(4):045015.

58. Bonham BH, Litvak LM. Current focusing and steering: modeling, physiology, and psychophysics. Hear Res. 2008;242(1-2):141–53.

